# Birth weight is not causally associated with adult asthma: results from instrumental variable analyses

**DOI:** 10.1101/471425

**Authors:** Ping Zeng, Xinghao Yu, Xiang Zhou

## Abstract

The association between lower birth weight and childhood asthma is well established by observational studies. However, it remains unclear whether the influence of lower birth weight on asthma can persist into adulthood. Here, we conducted a Mendelian randomization analysis to assess the causal relationship of birth weight on the risk of adult asthma. Specifically, we carefully selected genetic instruments based on summary statistics obtained from large-scale genome-wide association meta-analyses of birth weight (up to ~160,000 individuals) and adult asthma (up to ~62,000 individuals). We performed Mendelian randomization using two separate approaches: a genetic risk score approach and a two-sample inverse-variance weighted (IVW) approach. With 37 genetic instruments for birth weight, we estimated the causal effect per one standard deviation (SD) change of birth weight to be an odds ratio (OR) of 1.00 (95% CI 0.98~1.03, *p*=0.737) using the genetic risk score method. We did not observe nonlinear relationship or gender difference for the estimated causal effect. In addition, with the IVW method, we estimated the causal effect of birth weight on adult asthma was observed (OR=1.02, 95% CI 0.84~1.24, *p*=0.813). Additionally, the iMAP method provides no additional genome-wide evidence supporting the causal effects of birth weight on adult asthma. The result of the IVW method was robust against various sensitivity analyses, and MR-PRESSO and the Egger regression showed that no instrument outliers and no horizontal pleiotropy were likely to bias the results. Overall, this Mendelian randomization study provides no evidence for the fetal origins of diseases hypothesis for adult asthma, implying that the impact of birth weight on asthma in years of children and adolescents does not persist into adult and previous findings may be biased by confounders.

## Introduction

Asthma is a commonly complex chronic lung disease that is characterized by bronchoconstriction, airway hyper-responsiveness, mucus secretion and chronic inflammation (Wenzel 2012). Asthma represents a growing severe public health burden, affecting more than 300 million people and causing approximately 250,000 deaths per year worldwide (Olin and Wechsler 2014; Xu et al. 2014). Although asthma is very common in childhood, it can also occur in adulthood; for example, the incidence of asthma among adults is estimated to be as high as 12 cases per 1,000 person-years (Eagan et al. 2005; Masoli et al. 2004). Childhood and adult asthma share the same disease symptoms but likely have different genetic and environmental causes (Bush and Menzies-Gow 2009; Cabana et al. 2014; de Nijs et al. 2013; Larsen 2000; Ségala et al. 2008; Zihlif et al. 2016). In the literature it has been identified that various environmental, familial, socioeconomic, lifestyle and genetic factors are associated with asthma (e.g. obesity, smoking, air pollution and allergies) (Behrens 2009; de Nijs et al. 2013; Demenais et al. 2018; Ferreira et al. 2017; Polosa et al. 2008). Among these identified risk factors, the relationship between birth weight and subsequent risk of asthma has attracted much attention (Mu et al. 2014; Yuan et al. 2002). In practice birth weight is commonly employed as a proxy measurement of early life development and has long been hypothesized to have a profound long-term impact on individual’s predisposition to the risk of various diseases (e.g. asthma (Duijts 2012)) in later life — a hypothesis often referred to as the Barker hypothesis of adult diseases, or the fetal origins of adult diseases (Barker 1990; Barker 2004; Calkins and Devaskar 2011; Duijts 2012; Kindlund et al. 2010; Lucas et al. 1999). Indeed, it has been previously found in observational studies that lower birth weight is correlated with a higher risk of asthma in childhood and adolescence (Brooks et al. 2001; Carter et al. 2018; Kindlund et al. 2010; Liu et al. 2014; Seidman et al. 1991). Importantly, this inverse association between birth weight and childhood asthma is not confounded by familial factors (Örtqvist et al. 2009) and supported by large-scale meta-analyses (Lin et al. 2014; Mebrahtu et al. 2015; Mu et al. 2014; Xu et al. 2014).

However, whether lower birth weight still has a long-term influence on the risk of adult asthma is less understood and only very few studies have previously focused on this question (Johnson et al. 2015; Mu et al. 2014; Shaheen et al. 1999). It was estimated that the prevalence of asthma at 26 years among the lowest birth weight group (<2kg) was about twice higher that within the modal birth weight group (3~3.5kg) (Shaheen et al. 1999). Another result reported in (Johnson et al. 2015) implied that the asthmagenic effects of low birth weight can persist into adulthood, which was further supported by a recent meta-analysis where a 25% higher risk of adult asthma was observed among individuals with low birth weight (<2.5kg) compared with those with normal birth weight (2.5~4.0kg) (Mu et al. 2014). Nevertheless, it remains unclear whether the association between birth weight and adult asthma in these studies is truly causal as it is typical that other known/unknown factors (e.g. smoking or body mass index) in later childhood or early adulthood can confound the observed relationship between birth weight and adult asthma (Shaheen et al. 1999).

Understanding the long-term causal impact of birth weight on individual’s predisposition to asthma risk can facilitate our understanding of asthma etiology and paves ways for the potential development of early nutritional interventions to reduce asthma risk in adulthood. However, determining the causal impact of birth weight on adult asthma through traditional randomized intervention studies is challenging, as such studies necessarily require long-term follow-ups, are time-consuming, expensive, and often times, unethical (Eriksson et al. 2001; Lucas 1998). Therefore, it is desirable to determine the causal relationship between birth weight and adult asthma in observational studies using more novel statistical strategies (Sheehan et al. 2008). In the literature of causal inference, Mendelian randomization (MR) is a statistical tool that is commonly employed to determine the causal relationship between an exposure variable (e.g. birth weight) and an outcome variable (e.g. adult asthma) in observational studies. Specifically, MR is an instrumental variable method for causal inference that relies on associated single nucleotide polymorphisms (SNPs) from genome-wide association studies (GWASs) to serve as instruments (Angrist et al. 1996; Greenland 2000). By leveraging the fact that the two alleles of a genetic variant are randomly segregated during gamete formation and conception under the Mendel’s law and that such segregation is independent of various environmental confounders, MR analysis can provide estimate of causal effect without much susceptibility to reverse causation and other confounding factors as compared with other statistical approaches (Davey Smith and Ebrahim 2003).

In the present study we performed a MR study based on novel causal inference approaches including genetic risk score and two-sample inverse-variance weighted (IVW) estimation. Our study uses summary statistics obtained from large-scale GWASs with sample sizes ranging up to ~160,000 individuals for birth weight and ~62,000 individuals for adult asthma, representing the largest MR analysis performed to date for inferring the causal relationship between birth weight and adult asthma. Even with such large sample sizes, however, our study did not provide sufficient statistical evidence that supports the causal role of birth weight on adult asthma, suggesting that the previously detected association between birth weight and adult asthma may be unlikely a causal association.

## Materials and Methods

### Study design and data sources

We first obtained summary statistics in terms of marginal effect size estimates and their standard errors from the Early Growth Genetics (EGG) GWAS consortium study (Horikoshi et al. 2016). The EGG study is the largest GWAS formally published and performed to date on birth weight, which meta-analyzed a total of 16,245,523 genotyped and imputed SNPs on up to 143,677 individuals of European ancestry. In the EGG study, an additive linear regression model was applied to analyse one SNP at a time to detect the SNP associations with birth weight while properly controlling for gestational week and study-specific ancestry whenever these covariates were available (Horikoshi et al. 2016). With the EGG GWAS summary statistics, we yielded a set of 59 independent index SNPs that are strongly associated with offspring birth weight at the genome-wide significance level (*p*<5.00E-8) to serve as instruments (see Extended Data Table 1 shown in (Horikoshi et al. 2016) for full information).

A potential confounder for the birth-weight related MR analysis is the maternal effect — the portion of mother’s genetic effect on birth weight that is mediated through various maternal behaviors during pregnancy or intrauterine environment (Beaumont et al. 2018). To control for confounding due to the maternal effect, we first excluded instruments that exhibit potential maternal effects on birth weight using summary statistics from a recently published GWAS of maternal SNP effects on offspring birth weight (Beaumont et al. 2018). This maternal GWAS study collected 86,577 women and analyzed a total of 8,741,106 genotyped and imputed SNPs. While the sample size in the maternal GWAS is large, it is about half smaller compared with the EGG GWAS (86,577 vs. 143,702). Therefore, to effectively exclude all SNPs that may display observable maternal effects, we obtained a list of birth weight associated maternal SNPs based on a relaxed significant threshold (1.00E-5). In total, we obtained 700 candidate SNPs that may exhibit potential maternal effects. Afterwards, we then cross-examined the 59 instruments selected above with these 700 maternal SNPs and removed instruments that reside within 1Mb of any of the maternal SNPs. By this way, we excluded 12 instruments.

To exclude the potential pleiotropic effects, we also removed instrumental variables that are associated with relevant allergic diseases including asthma, hay fever and eczema. Specifically, we obtained GWAS summary results for these three allergic diseases from a recently published study (Demenais et al. 2018; Ferreira et al. 2017) and yielded the corresponding p values of the selected instruments for each disease. We then removed instruments that may show potential associations with asthma, hay fever or eczema (defined as *p*<0.05/58=8.62E-4). Excluding instruments that are strongly correlated to the outcome of interest (or outcome relevant traits) is a conservative strategy to guarantee the validity of the MR analysis — by focusing on only instruments that do not have horizontal pleiotropic effects, we can ensure that these instruments only have an influence adult asthma by the path of birth weight (Censin et al. 2017; Nelson et al. 2015; Østergaard et al. 2015). Afterwards, two additional instruments in this filtering step were excluded. We focused our following analysis on the remaining 45 instruments that unlikely exhibit maternal effects and unlikely exhibit pleiotropic effects.

Next, we obtained GWAS data from the Genetic Epidemiology Research on Aging (GERA) cohort (Banda et al. 2015). The GEAR study includes adult individuals whose age ranges from 18 to over 100 years old, with an average age of 63 years at the time of the survey in 2007, implying all the individuals including in our analyses are adult. After proper quality control [Hardy-Weinberg equilibrium (HWE) test *p* value<10^−4^, genotype call rate<95% and minor allele frequency (MAF) <0.01], we obtained a total of 487,609 SNPs on 61,916 people (24,718 males and 37,198 females; 10,101 of them (16.3%) have asthma). We phased genotypes using SHAPEIT (Delaneau et al. 2013) and imputed SNPs based on the Haplotype Reference Consortium (HRC version r1.1) reference panel (McCarthy et al. 2016) on the Michigan Imputation Server using Minimac3 (Das et al. 2016). After filtering (HWE *p* value<10^−4^, genotype call rate<95%, MAF<0.01 and imputation score<0.30), we obtained 8,385,867 genotyped and imputed SNPs. For each SNP in turn, we obtained summary statistics results (the effect size and its standard error) by using an additive logistic regression model while controlling for other available covariates (e.g. family income, education level, and gender, alcohol and smoking statuses, BMI and general health status as well as top ten principal components; descriptions of these covariates are shown in Table 1). Note that, among the set of 45 instruments for birth weight, only 37 are available in the GERA cohort. Therefore, we focused on these instruments in our MR analyses. For each of the remaining instruments in turn, we obtained summary statistics for both birth weight and adult asthma in terms of effect allele, marginal effect size, standard error and p value (Table 2).

**Table 1.** Descriptions of covariates available from the GERA cohort study

| Covariates | Code and proportion (%) |
| --- | --- |
| Education | 0: Other (4.97); 1: Elementary, High School or Technical School (12.6); 2: Some college (23.3); 3: College/Graduate School (59.1) |
| Income | 1: <\$39,999 (16.0); 2: \$40,000-\$59,999 (16.6); 3: \$60,000-\$99,999 (30.3); 4: \$100,000+ (37.1) |
| BMI | 1: ≤18 (1.66); 2: 19-25 (44.6); 3: 26-29 (28.7); 4: 30-39 (22.0); 5: 40+ (3.00) |
| Gender | 1: male (39.9); 2: female (60.1) |
| General health | 1: Excellent (19.3); 2: Very Good (38.2); 3: Good (33.7); 4: Fair/Poor (8.74) |
| Smoking | 0: 0 pack years (57.2); 1: ≤ 10 pack years (14.9); 2: 10~20 pack years (15.3); 3: 20~30 pack years (8.76); 4: >30 pack years (3.85) |
| Alcohol | 1: no days (35.6); 2: 1 day (14.3); 3: 2-4 days (20.8); 4: 5-6 days (10.5); 5: every day (18.8) |

Note: GERA: Genetic Epidemiology Research on Aging; BMI: body mass index; all the covariates were incorporated into the logistic regression as continuous variable.

**Table 2.** Summary statistics information for the selected instruments of birth weight and adult asthma

| Chr | SNP | Position | A<sub>1</sub>/A<sub>2</sub> | birth weight |  | adult asthma |  |
| --- | --- | --- | --- | --- | --- | --- | --- |
|  |  |  |  | beta | se | beta | se |
| 1 | rs2473248 | 22,536,643 | C/T | 0.0325 | 0.0057 | 0.0050 | 0.0224 |
| 1 | rs3753639 | 154,986,091 | C/T | 0.0306 | 0.0045 | -0.0175 | 0.018 |
| 1 | rs72480273 | 161,644,871 | C/A` | 0.0313 | 0.0051 | 0.0090 | 0.0205 |
| 1 | rs61830764 | 212,289,976 | A/G | 0.022 | 0.004 | 0.0020 | 0.0147 |
| 2 | rs1374204 | 46,484,205 | T/C | 0.047 | 0.0042 | -0.0326 | 0.0169 |
| 3 | rs11719201 | 123,068,744 | T/C | 0.0463 | 0.0044 | 0.0109 | 0.0181 |
| 3 | rs10935733 | 148,622,968 | T/C | 0.0221 | 0.0039 | -0.0067 | 0.0158 |
| 4 | rs925098 | 17,919,811 | G/A | 0.034 | 0.0042 | -0.0079 | 0.0175 |
| 5 | rs854037 | 57,091,783 | A/G | 0.0268 | 0.0048 | -0.0045 | 0.0201 |
| 5 | rs7729301 | 157,886,953 | A/G | 0.0239 | 0.0042 | -0.0068 | 0.0175 |
| 6 | rs35261542 | 20,675,792 | C/A | 0.0444 | 0.0041 | 0.0218 | 0.0173 |
| 6 | rs7742369 | 34,165,721 | G/A | 0.0283 | 0.0049 | -0.0075 | 0.0201 |
| 7 | rs798489 | 2,801,803 | C/T | 0.0233 | 0.0042 | -0.0254 | 0.0173 |
| 7 | rs6959887 | 35,295,365 | A/G | 0.0228 | 0.0038 | 0.007 | 0.015 |
| 8 | rs13266210 | 41,533,514 | A/G | 0.0308 | 0.0045 | 0.0237 | 0.0183 |
| 8 | rs6989280 | 126,508,746 | G/A | 0.0218 | 0.0042 | 0.0139 | 0.0175 |
| 8 | rs12543725 | 142,247,979 | G/A | 0.0231 | 0.0038 | -0.0172 | 0.0157 |
| 9 | rs28510415 | 98,245,026 | G/A | 0.0557 | 0.0065 | -0.0337 | 0.0269 |
| 9 | rs2150052 | 113,945,067 | T/A | 0.0211 | 0.0038 | 0.0109 | 0.0151 |
| 9 | rs700059 | 125,824,055 | G/A | 0.0334 | 0.0054 | 0.008 | 0.0243 |
| 10 | rs61862780 | 94,468,643 | T/C | 0.0281 | 0.0037 | -0.0274 | 0.0153 |
| 10 | rs74233809 | 104,913,940 | C/T | 0.0366 | 0.0067 | -0.0016 | 0.0262 |
| 10 | rs2421016 | 124,167,512 | T/C | 0.0207 | 0.0037 | 0.008 | 0.0154 |
| 11 | rs72851023 | 2,130,620 | T/C | 0.0476 | 0.0075 | -0.0491 | 0.0338 |
| 12 | rs12823128 | 26,872,730 | T/C | 0.0211 | 0.0037 | 0.0188 | 0.0155 |
| 12 | rs1351394 | 66,351,826 | T/C | 0.0436 | 0.0037 | -0.0151 | 0.0154 |
| 12 | rs7964361 | 102,994,878 | A/G | 0.0391 | 0.0067 | -0.0186 | 0.0274 |
| 13 | rs2324499 | 40,662,001 | G/C | 0.0217 | 0.004 | 0.0178 | 0.016 |
| 13 | rs2854355 | 48,882,363 | G/A | 0.0234 | 0.0044 | 0.0119 | 0.0171 |
| 13 | rs1819436 | 78,580,283 | C/T | 0.0329 | 0.0057 | -0.0082 | 0.0229 |
| 16 | rs1011939 | 19,992,996 | G/A | 0.0217 | 0.0041 | -0.0411 | 0.0172 |
| 17 | rs113086489 | 7,171,356 | T/C | 0.0307 | 0.0038 | -0.0479 | 0.0155 |
| 19 | rs10402712 | 33,926,013 | A/G | 0.0215 | 0.0043 | 0.0218 | 0.0176 |
| 20 | rs6040076 | 10,658,882 | C/G | 0.0231 | 0.0039 | -0.0215 | 0.0154 |
| 20 | rs6016377 | 39,172,728 | T/C | 0.0239 | 0.0039 | 0.006 | 0.0148 |
| 21 | rs2229742 | 16,339,172 | G/C | 0.036 | 0.006 | 0.0276 | 0.0254 |
| 22 | rs134594 | 29,468,456 | C/T | 0.0227 | 0.004 | 0.0257 | 0.0161 |

Note: Chr represents the chromosome; A<sub>1</sub> is the effect allele and A<sub>2</sub> is the alternative allele.

### Genetic risk score method

The genetic risk score (GRS) for birth weight was computed following (Guo et al. 2016; Ripatti et al. 2010). Briefly, the GRS for individual *i* in the GERA study was constructed as

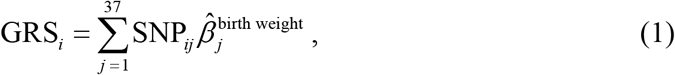

where 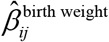 is the estimated marginal SNP effect size of birth weight for the *j*th instruments obtained from the EGG study (Horikoshi et al. 2016), and SNP_*ij*_ is the individual-level genotype of the effect allele for the corresponding *j*th instrument in the GERA study (Banda et al. 2015). We further standardized GRS to have mean zero and variance one in our analysis. Note that, unlike in (Guo et al. 2016; Ripatti et al. 2010) we did not scale GRS as the p value of GRS would not change regardless GRS was scaled or not. Afterwards, we evaluated the effect of GRS on adult asthma with an additive logistic regression model while controlling for family income, education level, and gender, alcohol and smoking statuses, BMI and general health status as well as top ten genotype principal components

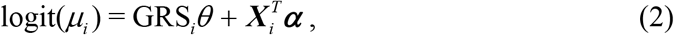

where *μ_i_* is the expectation of *y_i_* with *y_i_* = 1 and 0 representing the status of adult individual *i* with or without asthma in the GERA study, *θ* is the effect size of GRS, and ***X***_*i*_ is the vector of covariates with effect sizes ***α***. We are primarily interested in estimating *θ* and testing for the null hypothesis *H*_0_: *θ* = 0.

### Two-sample MR analysis

Besides the genetic risk score method, we also performed a two-sample MR analysis to estimate the causal effect size of birth weight on adult asthma using summary statistics data generated from the two consortia above. Suppose that the effect size estimate and its variance for the *j*th instrumental variable of birth weight are 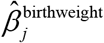 and 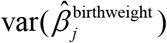 (*j* = 1, 2, …, 37), which are both obtained from the EGG study (Horikoshi et al. 2016). Suppose 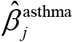 and 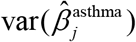 are the effect size estimate and its variance for the same instrumental variable in the GERA study (Banda et al. 2015), respectively. We estimated the causal effect of birth weight (again, denoted as *θ*) using all the instrumental variables together through the IVW method (Brockwell and Gordon 2001; Burgess et al. 2017; DerSimonian and Laird 1986; Hartwig et al. 2017; Thompson and Sharp 1999; Yavorska and Burgess 2017)

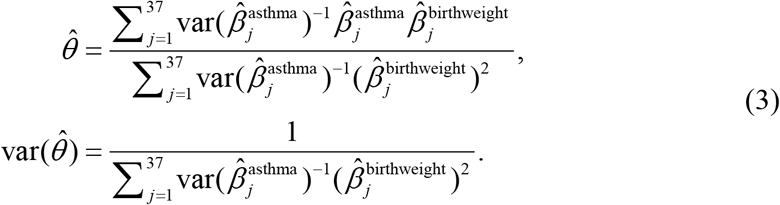

### The iMAP analysis to infer the causal effects

We further applied a recently developed method, iMAP, to complementally analyze the relationship between birth weight and adult asthma. iMAP is an integrative method for modeling pleiotropy and can be employed to investigate causality between pairs of complex traits using summary statistics from GWAS (Zeng et al. 2018). Unlike the genetic score or MR method, iMAP jointly analyzes all genome-wide SNPs and has the potential to provide additional evidence supporting or against causal relationship between pairs of traits. iMAP aims to estimate some proportional parameters that characterize the SNP causal effects on the two traits in order to better understand the relationship between the traits (Zeng et al. 2018). In particular, iMAP estimates an important ratio quantity π_11_/(π_10_+π_11_) (or π_11_/(π_01_+π_11_)), where *π_11_* represents the probability that a SNP is associated with both traits; π_10_ represents the probability that a SNP is associated with the first trait but not the second; π_01_ represents the probability that a SNP is associated with the second trait but not the first; and *π*_00_ represents the probability that a SNP is not associated with any traits. This estimated quantity above represents the proportion of SNPs associated with one trait that are also associated with the other and has been used to evaluate the causality of one trait on the other (Pickrell et al. 2016). Specifically, a large π_11_/(π_10_+π_11_) and a small π_11_/(π_01_+π_11_) imply that a large fraction of SNPs associated with the first trait is also associated with the second trait, but not vice versa, indicating that the first trait may causally affect the second trait. A small π_11_/(π_10_+π_11_) and a large π_11_/(π_01_+π_11_) indicate that the second trait may causally affect the first trait. On the other hand, a large π_11_/(π_10_+π_11_) and a large π_11_/(π_01_+π_11_) indicate that both traits may share common biological pathways. Therefore, estimating π_11_/(π_10_+π_11_) and π_11_/(π_01_+π_11_) using iMAP can help provide additional evidence with regard to the causal relationship between birth weight and adult asthma.

### Sensitivity analyses

To ensure results robustness and to guard against various modeling misspecifications in our main Mendelian mediation analyses, we performed extensive selectivity analyses. For the genetic risk score approach, we carried out stratified analysis in terms of gender. For the two-sample MR IVW method, we conducted the Mendelian randomization pleiotropy residual sum and outlier (MR-PRESSO) method to identify instrumental outliers that can substantially influence the causal effect estimate (Verbanck et al. 2018). In addition, we also conducted weighted median-based method which is robust when some instruments are invalid (Bowden et al. 2016a) as well as MR-Egger regression which guards against horizontal pleiotropic effects (Bowden et al. 2016b; Burgess and Thompson 2017).

### Power calculations

Finally, to investigate the statistical power, we carried out power calculations to detect a non-zero causal effect for birth weight (Brion et al. 2013; Burgess 2014; Freeman et al. 2013). In the calculations, we set the total phenotypic variance explained (PVE) by all instrumental variables to be 1.23% (i.e. the total phenotypic variance of birth weight explained by all used instruments; see below), set the significance level *α* to be 0.05, and set the proportion of the asthma cases equal to 16.3% (i.e. the fraction of cases observed in the GEAR study). In the present study, the powers were calculated using the method shown in (Brion et al. 2013) that can be easy to implement online at https://cnsgenomics.shinyapps.io/mRnd/.

## Results

### Estimated causal effect of birth weight on asthma with genetic risk score

We selected a set of 37 SNPs from a large-scale GWAS with up to 143,677 European individuals to serve as valid instruments for offspring birth weight (Table 1). These SNPs are all robustly associated with birth weight (*p*<5.00E-8) (Horikoshi et al. 2016), and explain a total of 1.23% phenotypic variance of birth weight based on summary statistics. We first examined the strength of these instrumental variables using *F* statistic (Noyce et al. 2017). The *F* statistics for all 37 SNPs are above 10 (ranging from 25.0 to 138.9 with mean 46.7), suggesting that all instruments are strong instruments and that weak instrument bias unlikely occurs in our analysis.

Using the logistic regression, we find that no causal association between the genetically determined birth weight and adult asthma after adjusting for available covariates. Specifically, the odds ratio (OR) per 1 standard deviation (SD) change of GRS is estimated to be 1.00, 95%CI 0.98~1.03, *p*=0.737. While the unadjusted OR for GRS is estimated to be 1.00, 95%CI 0.98~1.02, *p*=0.816. In addition, there is no evidence for quadratic effect size of GRS (*p*=0.602). We further implemented stratified logistic analysis for GRS separately in men and women. For men, the OR of per 1 SD change of GRS is estimated to be 0.97, 95%CI 0.93~1.01, *p*=0.108. For women, the OR of per 1 SD change of GRS is estimated to be 1.02, 95%CI 0.99~1.05, *p*=0.118. No quadratic effect of GRS on adult asthma is detected in men (*p*=0.846) or women (*p*=0.436).

### Estimated causal effect of birth weight on asthma with two-sample IVW method

In terms of the two-sample IVW method, the heterogeneity of casual effect of individual instrument is not observed (*p*=0.078) and the OR of per 1 SD change of offspring birth weight on adult asthma is estimated to be 1.02, 95%CI 0.84~1.24, *p*=0.813. Again, the results imply that there is no causal association between birth weight and adult asthma. MR-PRESSO shows that no outliers can substantially influence the estimate (*p*=0.113); see also Fig. 1A in which the SNP effect size of birth weight were plotted against the SNP effect size of adult asthma for each instrument. The weighted median method shows consistent null estimate (OR=0.91, 95%CI 0.67~1.23, *p*=0.533) and the Egger regression also generates similar null estimate (OR=0.78, 95%CI 0.36~1.72, *p*=0.540). The intercept of the Egger regression is not significantly deviated from zero and is estimated to be 0.008 (95% CI −0.015~0.032, *p*=0.483), suggesting that the assumption of balanced pleiotropy holds in our two-sample MR analysis. The funnel plot for individual causal effect size estimated for each single instrument also demonstrates a symmetric pattern of effect size variation around the point estimate (Fig. 1B). Together, MR-Egger regression intercept and funnel plot indicate that horizontal pleiotropy unlikely bias our results.

**Fig. 1.**
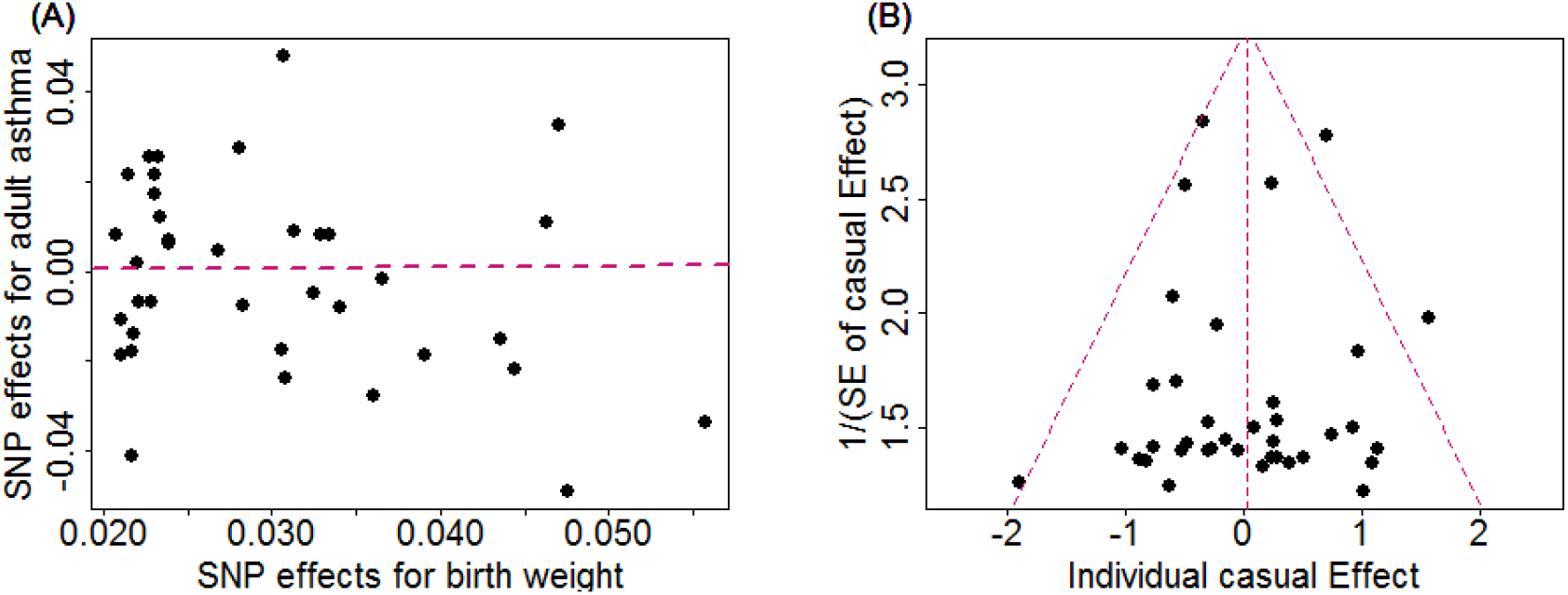
(**A**) Relationship between the SNP effect size estimates of birth weight (x-axis) and the corresponding effect size estimates of adult asthma (y-axis) using 37 instruments. The line in red represents the estimated casual effect of birth weight on adult asthma obtained using the IVW method. (**B**) Funnel plot for single causal effect estimate of birth weight on adult asthma. The vertical line in red represents the estimated casual effect of birth weight on adult asthma obtained using the IVW method

### Results of the iMAP method

Using the iMAP method (Zeng et al. 2018), the proportion of SNPs associated with birth weight which is also associated with adult asthma is estimated to be 7.43E-4, the proportion of SNPs associated with adult asthma that is also associated with birth weight is 6.59E-5. Both the proportions are rather small, suggesting that SNPs associated with the birth weight are unlikely to be associated with adult asthma. The result of iMAP above is not a surprising as it is shown that no association signals are overlapped between birth weight and adult asthma (Fig. 2). Additionally, the overall genetic correlation was only estimated to be 0.050 (se=0.069, *p*=0.471) using the linkage disequilibrium score regression (LDSC) (Bulik-Sullivan et al. 2015). Therefore, iMAP provides no additional genome-wide evidence supporting the causal effects of birth weight on adult asthma.

**Fig. 2.**
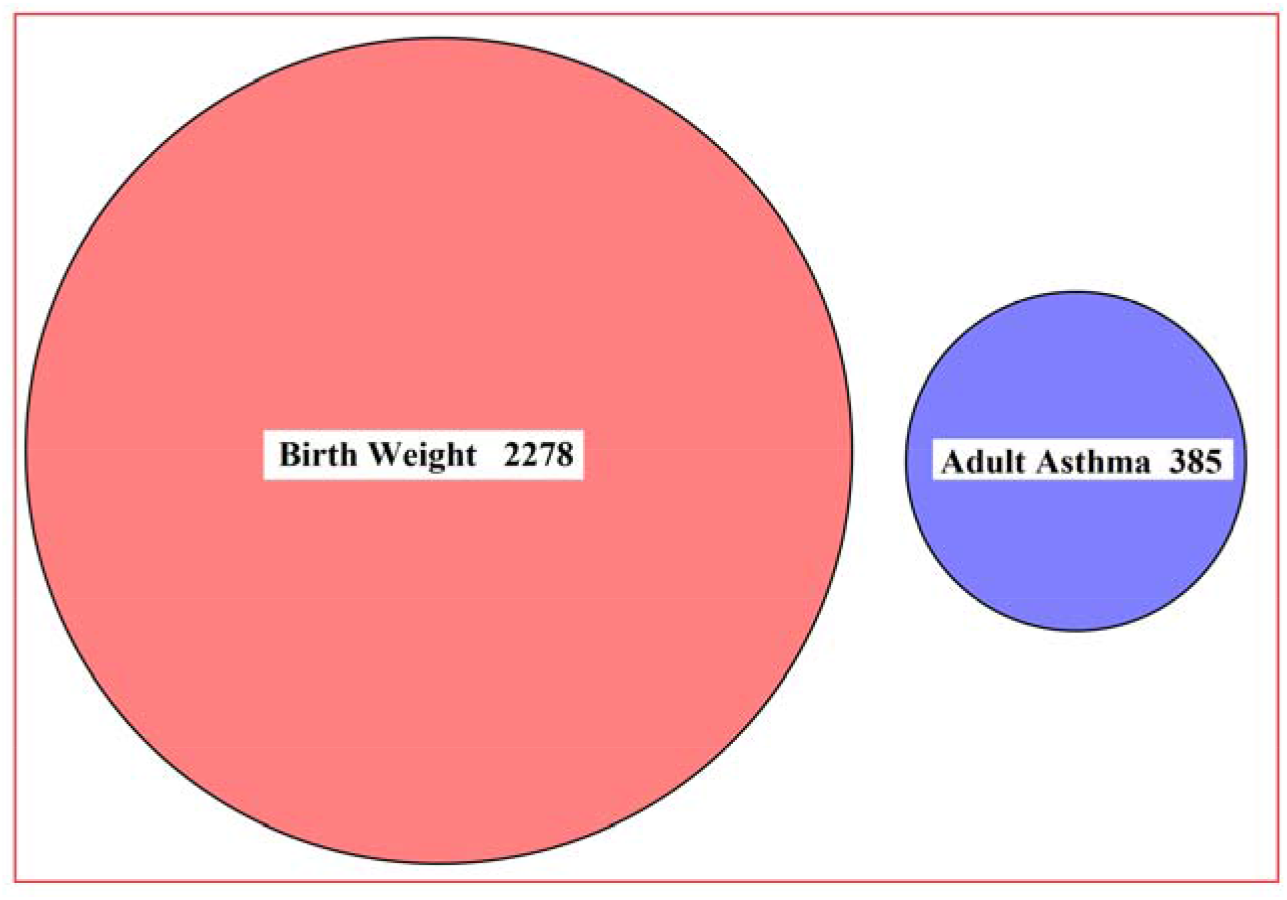
Venn diagram displays the association pattern of genome-wide significant SNPs (*p*<5E-8) shared by birth weight and adult asthma. There are 2,278 and 385 identified SNPs associated with birth weight and adult asthma, respectively; while no significant SNPs shared by birth weight and adult asthma

### Results of power calculations

We finally examine whether the lack of detectable non-zero causal effect of birth weight on adult asthma is due to a lack of statistical power. To do so, we performed the statistical power calculations to detect an OR ratio of 1.10, 1.2 or 1.3 in the risk of adult asthma per unit change of birth weight following the approach shown in (Brion et al. 2013). Note that the effect of birth weight on adult asthma was previously estimated to be 1.25 (Mu et al. 2014). It is shown that the estimated statistical power is 17%, 51% and 84%, respectively, implying that we would have moderate or high power to detect such a causal effect of birth weight on adult asthma.

## Discussion

In the present paper, we have explored the fetal origins of adult asthma hypothesis by performing a comprehensive Mendelian randomization analysis to investigate the causal effects of birth weight on adult asthma. To efficiently avoid possible violation of model assumptions, we have carefully chosen SNPs to serve as valid instruments and conducted extensive sensitivity analyses to ensure the validity of Mendelian randomization analysis (Fritsche et al. 2016; Noyce et al. 2017). With valid instruments from large scale GWAS of birth weight we have demonstrated that the genetically increased/decreased birth weight is not casually associated with adult asthma.

Our results are not consistent with the previous associations between birth weight and asthma discovered in observational studies. However, the associations between birth weight and adult asthma in these previous observation studies may be confounded by many known/unknown confounders that occur during prenatal or postnatal life (e.g. the adult body mass index, BMI, and smoking status in adulthood) (Carter et al. 2018; Shaheen et al. 1999). Therefore, the association previously detected in observational studies could be spurious associations. Indeed, by using a propensity score approach to control for confounders, it has been showed that birth weight is not associated with the risk of asthma during the first 6 years of life (Yang et al. 2013). In addition, after considering the maternal smoking status in pregnancy (Carter et al. 2018) and gestational age (Liu et al. 2014; Sonnenschein-van der Voort et al. 2014), the estimated association size between lower birth weight with asthma is much reduced. Therefore, our GRS approach and MR analyses are consistent with these observational studies that properly control for confounding effects, providing additional evidence suggesting that birth weight may not be causally associated with adult asthma.

Finally, we emphasize that we cannot completely rule out the possibility that we are underpowered to discover a weak causal influence of birth weight on adult asthma as shown in the power calculations. Completely classify this issue requires dataset of adult asthma with larger sample size in the future. We also note that two large scale GWASs about adult asthma were published recently and the corresponding summary statistics results can be publicly available (Demenais et al. 2018; Ferreira et al. 2017).

However, due to the following reasons, we did not consider either of these two datasets. In particular, in the study of Ferreira et al (Ferreira et al. 2017), the analysis was performed on three allergic diseases (i.e. asthma, hay fever and eczema elucidates), thus the asthma-specific summary statistics results cannot be obtained. Additionally, about half samples in the birth weight EGG study and about 40% individuals in Ferreira et al (Ferreira et al. 2017) come from the same UK BioBank data resource (Sudlow et al. 2015), leading to the issue of sample overlapping in the MR analysis. Sample overlapping can result in severely biased causal effect estimates and adjusting for sample overlapping is statistically challenging (Burgess et al. 2016). For the study of Demenais et al (Demenais et al. 2018), the summary statistics of adult asthma are available for only 16 instruments (vs 37 in the GERA study). The lack of instruments can result in a substantial loss of information and potentially lead to weak instruments bias. Indeed, using 16 instruments and summary statistics from Demenais et al (Demenais et al. 2018), we obtained a similar estimate of causal effect for birth weight on adult asthma (OR=1.05, 95%CI 0.82~1.34, *p*=0.701), again supporting our conclusion above.

## Conclusions

Overall, our results do not provide support for the fetal origins of diseases hypothesis for adult asthma, implying that the impact of birth weight on asthma is less possible to last into adult and some of the previous findings on the association between birth weight and asthma may be biased by confounders.

## Acknowledgements

We thank all the GWAS consortium studies for making the summary data publicly available and are grateful of all the investigators and participants contributed to those studies. The GERA Data came from a grant, the Resource for Genetic Epidemiology Research in Adult Health and Aging (RC2 AG033067; Schaefer and Risch, PIs) awarded to the Kaiser Permanente Research Program on Genes, Environment, and Health (RPGEH) and the UCSF Institute for Human Genetics. The RPGEH was supported by grants from the Robert Wood Johnson Foundation, the Wayne and Gladys Valley Foundation, the Ellison Medical Foundation, Kaiser Permanente Northern California, and the Kaiser Permanente National and Northern California Community Benefit Programs. The RPGEH and the Resource for Genetic Epidemiology Research in Adult Health and Aging are described in the following publication, Schaefer C, et al., The Kaiser Permanente Research Program on Genes, Environment and Health: Development of a Research Resource in a Multi-Ethnic Health Plan with Electronic Medical Records, in preparation, 2013. This work was supported by the National Institutes of Health R01HG009124, the National Science Foundation DMS1712933, the National Natural Science Foundation of Jiangsu Province (BK20181472), the Youth Foundation of Humanity and Social Science funded by Ministry of Education of China (18YJC910002), the China Postdoctoral Science Foundation (2018M630607), Jiangsu QingLan Research Project for Outstanding Young Teachers, the Postdoctoral Science Foundation of Xuzhou Medical University, the National Natural Science Foundation of China (81402765), the Statistical Science Research Project from National Bureau of Statistics of China (2014LY112), and the Priority Academic Program Development of Jiangsu Higher Education Institutions (PAPD) for Xuzhou Medical University.

